# NeuroCSF: an fMRI method to measure contrast sensitivity function in human visual cortex

**DOI:** 10.1101/2024.01.22.576727

**Authors:** Laurie Goulet, Reza Farivar

**Affiliations:** Department of Ophthalmology & Visual Sciences, McGill University, Montreal, Canada

**Keywords:** Contrast sensitivity function, visual field, fMRI, model-driven approach, visual perception

## Abstract

The contrast sensitivity function (CSF) describes a range of spatial frequencies (SF) that are detectable at a given level of contrast and is a very valuable tool both in clinical and fundamental research. However, despite its immense value, the full potential of the CSF has not been utilized in every aspect of clinical research due to time limits and patient factors. We propose neuroCSF as a new method for measuring the CSF across the visual field directly from brain activity, and with minimal demand from participants. NeuroCSF is a computational model that estimates voxel-wise CSF parameters (i.e., peak contrast sensitivity, peak spatial frequency, and spatial frequency bandwidth) from functional magnetic resonance imaging (fMRI) signals, under controlled visual stimulation conditions. The approach extends the population spatial frequency tuning (Aghajari, Vinke, & Ling, 2020) and population receptive field (Dumoulin & Wandell, 2008) methods to provide the first characterization of a full CSF using neuroimaging. We observe that across early visual areas (V1, V2 and V3), the CSF peak spatial frequency and spatial frequency cutoff are significantly higher for foveal eccentricity and decrease at parafoveal eccentricities. Conversely, SF bandwidth slowly increases with eccentricity, while peak contrast sensitivity remains constant with eccentricity for all early visual areas. Thus, cortical CSF estimates vary systematically with eccentricity. The neuroCSF approach opens new perspectives for the study of cortical visual functions in various disorders where the CSF is impacted, such as amblyopia, traumatic brain injury, and multiple sclerosis.

## 1. Introduction

The Contrast Sensitivity Function (CSF), originally developed by Campbell and Robson (1968) describes the range of spatial frequencies detectable at varying levels of contrast. It is widely acknowledged as the gold standard for evaluating visual performance. The CSF not only serves to delineate the boundary between perceptually visible and invisible stimuli but also sheds light on the limiting factors affecting everyday functional vision (Gervais, Harvey, & Roberts, 1984; Vision, 1985).

The CSF predicts visual target recognition in everyday life, encompassing tasks such as recognizing faces (Owsley & Sloane, 1987; Stalin & Dalton, 2020; West et al., 2002), interpreting road signs (Owsley & Sloane, 1987), deciphering letters, and performing real-world visual tasks like reading, driving, and mobility (Legge, Rubin, Pelli, & Schleske, 1985; Lovie-Kitchin, Mainstone, Robinson, & Brown, 1990; Marron & Bailey, 1982; Woods & Wood, 1995). Its ability to assess the visual system’s sensitivity to different levels of contrast provides crucial information about the overall health and functioning of the system. In clinical visual sciences (Arden, 1978), the CSF holds a vital place, serving as a non-invasive assessment tool for the detection and tracking of visual impairments originating from ocular diseases or neurological deficits. Below, we briefly discuss the clinical concerns for which CSF can be valuable (amblyopia, age-related macular degeneration, and multiple sclerosis) and speculate on a fourth—post-concussive visual blur.

Amblyopia, commonly known as “lazy eye”, is characterized by reduced visual acuity in an otherwise healthy eye and is often resulting from abnormal visual experience in early childhood. It is traditionally diagnosed with visual acuity tests, but losses in contrast sensitivity have been detected in amblyopes with reported normal visual acuity (Jindra & Zemon, 1989). Studies report significant contrast sensitivity reductions in both the treated amblyopic eye (AE) and fellow-fixing eye (FE) of "cured" amblyopic subjects when compared to control subjects (Cascairo, Mazow, Holladay, & Prager, 1997; Chatzistefanou et al., 2005; Huang, Tao, Zhou, & Lu, 2007; Rogers, Bremer, & Leguire, 1987; Sjöstrand, 1981; Wang, Zhao, Ding, & Wang, 2017). Analyzing the CSF could thus aid in the detection and monitoring of amblyopia, providing insights into visual symptoms that go beyond acuity measurements.

Age-related macular degeneration (AMD) is a progressive retinal disease that primarily affects the central portion of the retina that is responsible for detailed vision. It is one of the leading causes of vision impairments in the elderly (Wong et al., 2014). Early changes of AMD are predominantly morphological and often occur without noticeable or measurable vision loss symptoms. Conventional clinical tests, such as visual acuity, often miss these early subtle changes, limiting their clinical utility in early and intermediate stages of AMD (Cocce et al., 2018; Klein, Wang, Klein, Moss, & Meuer, 1995; Owsley, Huisingh, Clark, Jackson, & McGwin, 2016). Measuring early functional changes in AMD poses significant challenges and the CSF could be instrumental in tracking functional vision loss associated with AMD (Hoffmann, Rossouw, Guichard, & Hatz, 2020). In its early stage, AMD results in a CSF that is similar in shape to the normal population, but that is shifted along both axes (Chung & Legge, 2016; Kleiner, Enger, Alexander, & Fine, 1988). As the disease progresses, the CSF can reveal specific patterns of deficits within the visual field, allowing clinicians to tailor interventions and monitor their effectiveness.

Multiple sclerosis (MS), a disorder involving demyelination of the central nervous system, is commonly manifested early on by ocular symptoms. Findings suggest that MS involves contrast sensitivity deficits that precede changes in visual acuity (Chen & Gordon, 2005; Jackson, Ong, & Ripley, 2004; McDonald & Barnes, 1992). A recent study by Nunes, Monteiro, and Vaz Pato (2014) revealed that in its early stages, MS can produce highly selective contrast sensitivity loss for high spatial frequencies. As the disease progresses, greater losses are seen in the low-frequencies band. By assessing the contrast sensitivity function, clinicians can identify early signs of MS in patients and track the progression of the disease.

While CSF degradation has been often considered in light of optical or ocular losses, in at least one case—post-traumatic visual blur—the eyes and the optics appear normal but patients report persistent “blur” (Collins et al., 2016; Ripley & Politzer, 2010). Because this type of blur cannot be readily explained by ocular or optical factors, it necessitates neurological measures of the CSF, which currently do not exist.

The major problem faced is that despite its immense value, the full potential of the CSF has not been utilized in every aspect of clinical research, due to several challenges associated with standard psychophysical CSF techniques, including time constraints, patient-related issues, and technique limitations.

One of the primary challenges comes from the substantial time required to perform a full CSF assessment at one retinal location (typically foveal), ranging from 15 to 30 minutes per eye. This duration that already poses as a challenge for healthy participants can become almost impossible to perform for a patient that also suffers from attention and/or fixation problems, visual discomfort, or mobility issues (Harvey, 1997; Kelly & Savoie, 1973). Behavioral CSF measures are also affected by extraneous factors such as fatigue, compliance, attention (Abramov et al., 1984; Bradley & Freeman, 1982), emotional arousal (Lee, Baek, Lu, & Mather, 2014) and feedback adaptation(Abrahamyan, Silva, Dakin, Carandini, & Gardner, 2016).

Crucially, CSF measures are typically done at one retinal location—foveally. In certain visual deficits like amblyopia or age-related macular degeneration, visual impairments vary *across* the visual field (Katz, Levi, & Bedell, 1984; Midena & Pilotto, 2017; Thomas, 1978). Currently to obtain information about such visual field distinctions, the CSF must be behaviorally tested at multiple positions, which poses challenges for both healthy and clinical participants. This is why the standard laboratory methods for measuring the CSF have not been used to their full potential in clinical research.

In recent years, various alternative approaches to psychophysical CSF measurement have been proposed, but often at the cost of efficiency or validity. For instance, Visual Evoked Potentials (VEPs) offer objectivity but vary in sensitivity (Hemptinne, Liu-Shuang, Yuksel, & Rossion, 2018; Howe & Mitchell, 1984; Lopes de Faria, Katsumi, Arai, & Hirose, 1998). Similarly, methods relying on eye movements, such as Gradiate (Mooney, Alam, Hill, & Prusky, 2020) or optokinetic nystagmus (OKN) (Dakin & Turnbull, 2016; Jones, Kalwarowsky, Atkinson, Braddick, & Nardini, 2014), are reliable approaches that can be used with children or individuals with cognitive deficits, but vary in complexity and efficacy, and fail to account for retinal location differences.

The CSF is a promising tool that can quantify subtle patterns of visual deficits that may otherwise go unrecognized by current testing methods and provides information relevant for disease progression and monitoring of treatment efficacy. Existing tools have a number of constraints, mainly demanding extensive behavioural compliance and limited assessment at multiple retinal locations. We sought to overcome these challenges to develop an objective measure of the CSF across the visual field with minimal behavioural demands on the patient.

We have developed the neuroCSF, a method for directly estimating the CSF across the visual field from functional Magnetic Resonance Imaging (fMRI) signals. The neuroCSF objectively estimates voxel-wise CSF parameters, including peak contrast sensitivity, peak spatial frequency, and spatial frequency bandwidth. This estimation process relies on a modeling approach that translates the behavioral CSF model into a neural one.

The neuroCSF offers a substantial advantage over traditional approaches by simultaneously measuring all visual field locations within an eccentricity limit in under 35 minutes of scan time (and potentially much shorter, with further optimizations). It provides unbiased measures of the CSF for each voxel across the visual brain without requiring input from participants. This transformative method represents a significant advancement in CSF measurement, offering an objective, fixation-independent, and non-invasive method to simultaneously assess the entire visual field.

## 2. Materials and Methods

### 2.1. Subjects

Eight subjects with normal or corrected-to-normal vision (3 males; mean age, 27; range, 22-32 years) were recruited for this experiment. The participants had no prior history of binocular dysfunction. Informed consent was obtained from each subject in accordance with the Code of Ethics of the World Medical Association (Declaration of Helsinki) and was approved by the Research Ethics Board of the McGill University Health Center.

### 2.2. Imaging

#### 2.2.1. Display

Stimuli were generated using Psychtoolbox (Brainard, 1997) through MATLAB (2019a, The Math Works Inc., Natick, Massachusetts) and displayed on a gamma-corrected 32” LCD BOLDScreen (Cambridge Research Systems) at 120Hz with a mean luminance of 120 cd/m^2^ reflected by a mirror above the participant’s head. Participants were placed at a viewing distance of 152 cm from the monitor spanning 25.8 by 14.7 degrees of visual angle at a pixel resolution of 1920 by 1080. All stimuli were viewed binocularly.

#### 2.2.2. Equipment/setup

Acquisition was performed on a Siemens Prisma 3T MRI Scanner at the Montreal General Hospital (McGill University Health Center). Functional images were acquired using a 32-channel vision coil (Farivar, Grigorov, van der Kouwe, Wald, & Keil, 2016) in a transverse plane perpendicular to the calcarine sulcus with a standard simultaneous multislice (SMS) echo-planar imaging (EPI) sequence (Resolution = 2 x 2 x 2 mm, TR= 1000ms, TE: 33ms, FA= 33°, fat saturation, FOV = 128 x 128 mm, number of slices= 16, R=2, BW = 1698 Hz/pixel). Anatomical images were obtained with a 64-channel head coil using a T1-weighted magnetization prepared-rapid gradient echo (MPRAGE) sequence (Resolution = 0.8mm3, TR= 1780ms, TE=2.64ms, FA = 9o, FOV = 216 x 216 mm, number of slices= 208).

#### 2.2.3. Preprocessing of Functional Images

Preprocessing of fMRI data was performed using Analysis of Functional NeuroImages (AFNI) (Cox, 1996). To minimize spatial blurring, all spatial transformations following slice-time correction (i.e., motion correction, distortion correction, registration to anatomical image) were applied in a single step. Preprocessing of the T1-weighted anatomical images for alignment with the functional images was performed using Freesurfer (http://surfer.nmr.mgh.harvard.edu/).

During the functional runs, we binocularly presented full-field stimuli that varied systematically in contrast (min=10 % max=60%) and spatial frequency (min=0.033 cyc/° [cpd], max=26cpd). Stimuli were generated by overlapping eight sinusoidal gratings with evenly spaced rotations over 180°, making the stimuli not selective for any orientation.

### 2.3. NeuroCSF Stimuli & Experimental Procedure

Each run included 20 ten-second periods of smoothly transitioning contrast-spatial frequency vectors—crossing from the subthreshold to suprathreshold regimes of the CSF (Fig. 1)—presented in a random sequence. The crossing points were evenly spaced (logarithmically) across the expected range in the contrast-spatial frequency space. This smoothly dynamic stimuli strategy ensured that we would capture activity, as measured through the blood oxygen level-dependent (BOLD) response via fMRI, as it transitioned from zero response—when the stimulus was subthreshold—to complete activation—when the stimulus was suprathreshold. We thus captured the experience of an entire transition from subthreshold to suprathreshold and observed the changes in neural activity as the brain began to detect the stimuli.

**Figure 1.**
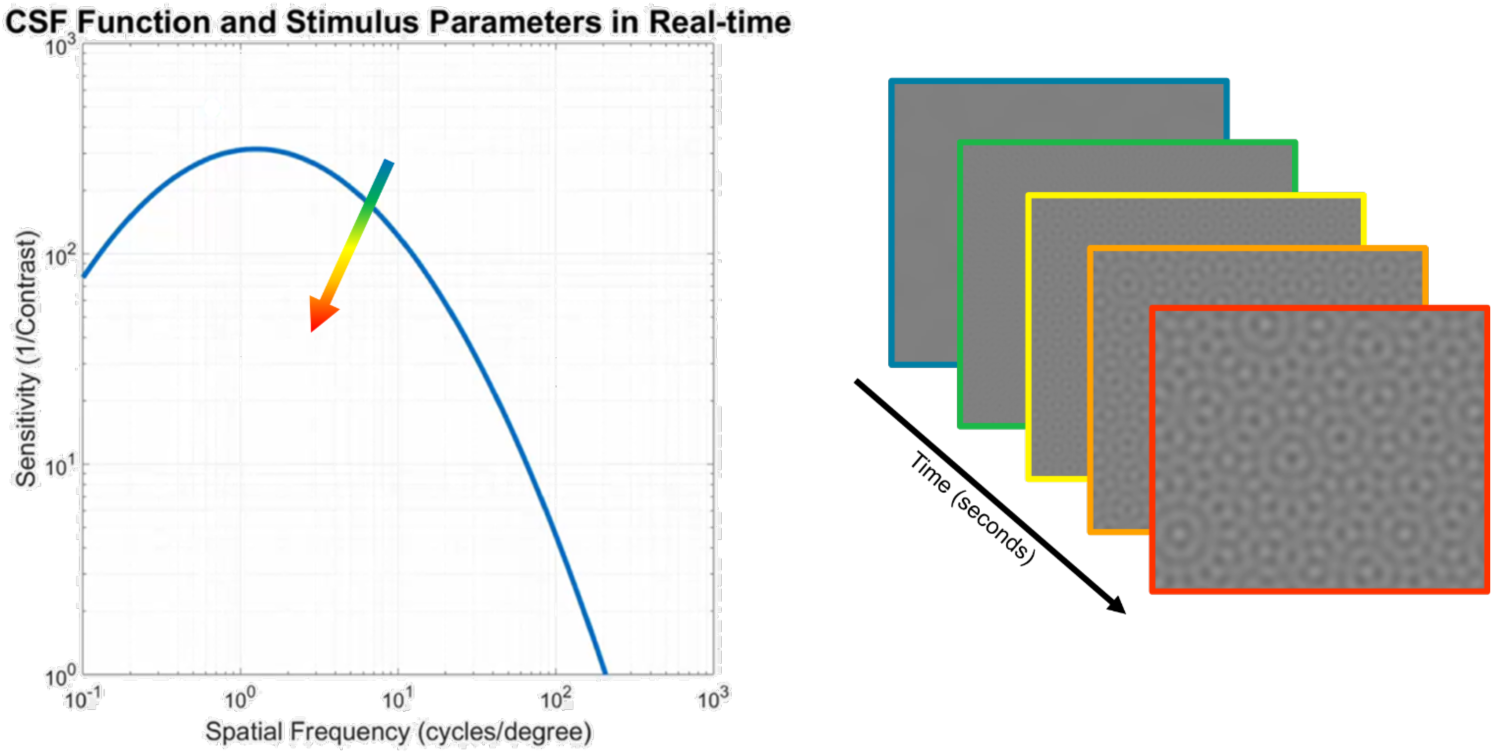
Stimulus presentation sequence for the CSF modeling. Depiction of one transition of the full field dynamic stimuli that varies over time in spatial frequency and contrast (right). The five frames depicted are only a subset of the 300 frames that would be presented within one of the 20 stimulus transitions. The stimulus consists of a contrast and spatial frequency vector (left) and any of its time points represent a shift in this two-dimensional space of contrast and spatial frequency.

Each stimulus presentation was followed by a blank interstimulus interval defined by a Gaussian distribution with a mean of 5s. To help with attention, participants were instructed to report whenever they perceived the stimuli by pressing a key. Each session comprised of three unique runs, with the same trajectories but shown in different orders, repeated twice, totaling approximately 30 minutes.

### 2.4. NeuroCSF Data Analysis

The different CSF transitions that we presented to the subjects during fMRI scan were recorded along with the contrast, spatial frequency, and the precise transition time (Fig. 2A). The challenge then becomes to identify the CSF model that, when taken into account to generate the model hemodynamic response given the stimulation pattern, explains the bulk of the fMRI response.

**Figure 2.**
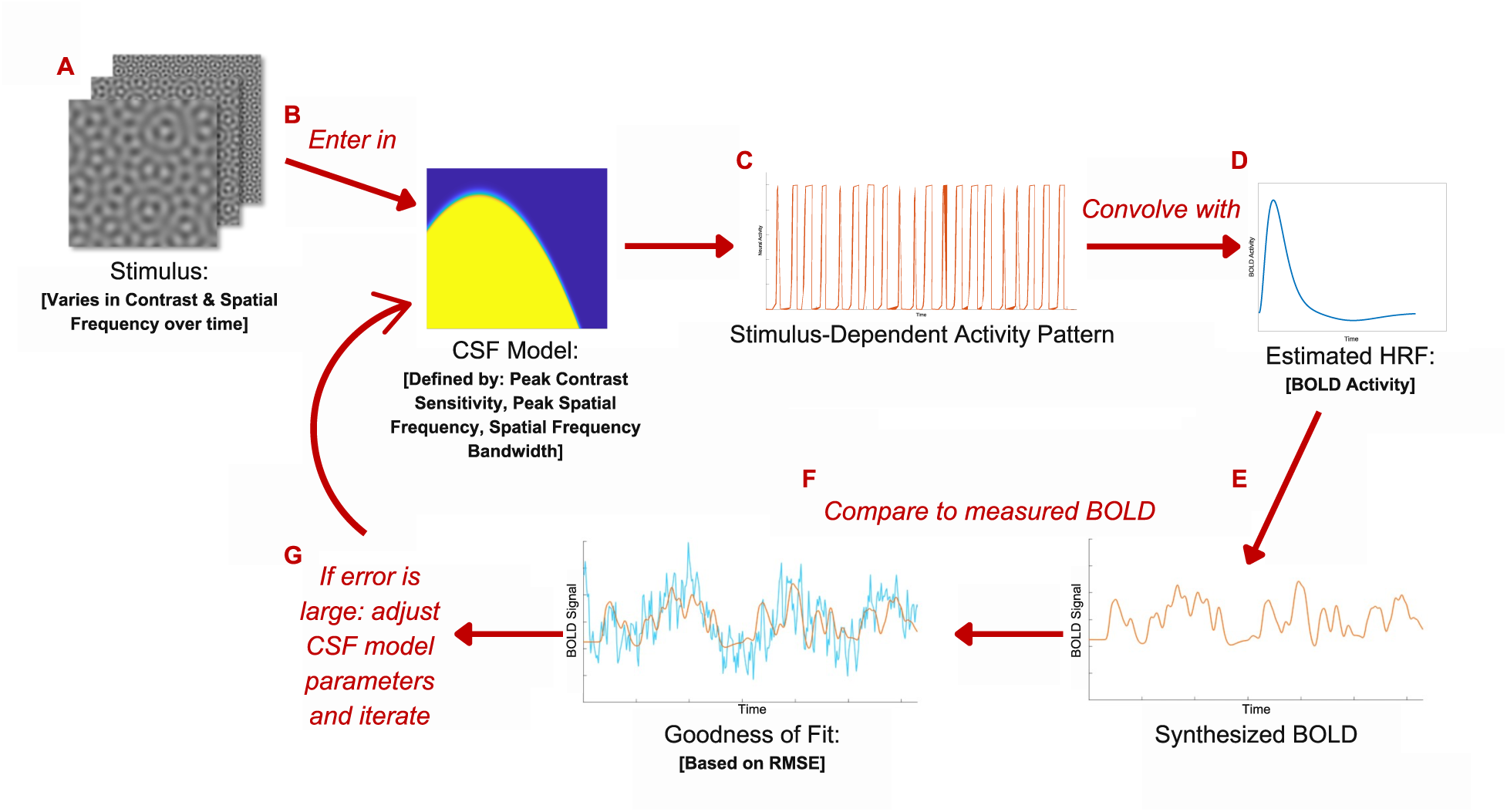
Depiction of the CSF model fitting method. The procedure begins by assigning a canonical CSF to a voxel (B) and, in consideration of the temporal profile of the stimulus (A), predicting the time-varying voxel response (C). This vector, convolved with the hemodynamic response (D), yields a synthesized fMRI timeseries model (E), which can then be fitted to the measured fMRI data from the voxel (F). If the fit does not sufficiently explain variability in the voxel’s time series, we iterate until we converge on a parameter set that best explains the voxel’s response (G).

The CSF model was estimated by assigning an initial model of the CSF to each voxel, defined by three parameters (i.e., peak contrast sensitivity, peak spatial frequency, spatial frequency bandwidth) (Fig. 2B). The mathematical model of the CSF can be represented by a log-parabola, S’(f), defining sensitivity as (Watson & Ahumada, 2005):

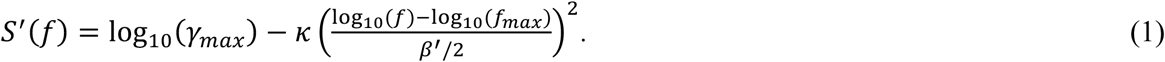

Where *κ* = log_10_(2), *β*^′^ = log_10_(2*β*), γ*_max_* represents the peak sensitivity (1/Contrast), *f_max_* the peak spatial frequency in cycles per degree (cpd) and β the spatial frequency bandwidth (octaves).

In this approach, the CSF model of each voxel was treated similarly to a receptive field representing a boundary in log-log space of contrast sensitivity and spatial frequency. Any time-point of our stimulus (Fig. 2A) represented a shift in this 2-D space of contrast and spatial frequency, such that when the stimulus entered a voxel’s receptive field or CSF boundary, and was now suprathreshold, this would elicit a response from the voxel. By calculating the overlap between the stimulus sequence (Fig. 2A) and the CSF model (Fig. 2B), we defined a time-varying voxel response (Fig. 2C). The time-varying voxel response vector was then convolved with a subject-specific and voxel-specific estimated hemodynamic response function (Fig. 2D) to produce a synthesized BOLD timeseries. This produced a function that defined proportional BOLD activity at every time point (Fig. 2E). This model BOLD time-series was then compared to the measured BOLD time-series (Fig. 2F). The difference between the two was calculated using a Goodness of Fit measurement (i.e., RMSE).

If the fit explained variability in the voxel sufficiently, the process stopped, and the parameters of the initial CSF model were saved for that voxel. If not, we searched a parameter space using a particle swarm optimization algorithm (Ab Wahab, Nefti-Meziani, & Atyabi, 2015; Helwig, Branke, & Mostaghim, 2013; Mendes, Kennedy, & Neves, 2004) and iterated until we converged on the set of CSF parameters that resulted in the best fit between the model and measured BOLD timeseries (Fig. 2G). This process was repeated for every voxel in areas V1, V2, and V3.

Based on the estimated CSF parameters (i.e., peak sensitivity, peak spatial frequency and spatial frequency bandwidth), we then derived voxels’ high spatial frequency cutoff from equation 1 by setting *S*^′^(*f*) to 1 and then calculated the associated spatial frequency.

### 2.5. Hemodynamic Response Function (HRF) Estimation

Since the shape of the hemodynamic response function (HRF) can vary significantly between people, we estimated individual HRF for each participant and modeled personalized HRF for each voxel within our region of interest (ROI).

#### 2.5.1. HRF-mapping stimulus

The stimulus consisted of a full field (radius 16◦ visual angle), high contrast texture, consisting of a dynamic, high-contrast pseudo-checkerboard varying in spatial frequency and phase function (Alvarez, De Haas, Clark, Rees, & Schwarzkopf, 2015), that briefly appeared on the display for two seconds before returning to a static grey screen for a 20 second period. This cycle was repeated 10 times, totalling one run of approximatively 5 minutes. Participants were instructed to maintain their gaze on a central fixation point (size: 0.1 degree), and to report the color change of the fixation dot.

#### 2.5.2. HRF modelling

The HRF was fitted with the difference of two gamma density functions (Worsley et al., 2002). The modeled parameters were the time-to-peak of the HRF response and HRF undershoot, the full width at half maximum of the HRF response and HRF undershoot, the ratio adjusting the amplitude of the HRF undershoot relative to the HRF response amplitude, and the scale factor for the final HRF. The HRF parameters were optimized simultaneously using particle swarm non-linear optimization algorithm.

### 2.6. Retinotopic Atlas ROIs

Surface location and retinotopy estimates (eccentricity and polar angle) of early visual areas (V1,V2, and V3) were obtained by registering a probabilistic retinotopic atlas (N. C. Benson et al., 2012) (https://cfn.upenn.edu/aguirre/wiki/doku.php?id=public:retinotopy_template) to each subject’s functional space.

### 2.7. NeuroCSF Voxel Selection

Only voxels with a receptive field center within the range of 0 to 6 degrees of eccentricity were included in the analysis. We also only included gray matter voxels whose coefficient of determination (R^2^) was greater than the highest R^2^ obtained in any white matter voxel (i.e., use the distribution of R^2^ in the white matter as a sample of trivial R^2^ values). Based on this method, only voxels with a minimum R^2^ of 10% were included in the analysis.

### 2.8. Cross-validation

Each subject was observed with fMRI while viewing three unique runs of our novel stimulus presentation. Each of these runs had a pair with perfectly matched presentation time, for a total of 6 runs. To maximize the R^2^ for the fit, we averaged the fMRI data from pairs of runs and thus obtained three measurements of CSF parameters for each voxel. To validate the CSF parameters, we employed leave-one-out-cross-validation (LOOCV) for each voxel in the early visual cortex. Parameters from two pairs of runs were used to predict the time series of another pair of runs, and an average CSF parameter set per voxel was calculated. Cutoff spatial frequency was then derived for each voxel from equation 1.

## 3. Results

3.1. Cortical CSF parameters consistency and eccentricity variability in early visual areas

We first qualitatively assessed the organization of the CSF parameters across the visuocortical map of early visual areas. Parameter estimates were projected onto the cortical surface (Fig. 3) and the visual field maps (Fig. 4-7) to visualize local variations in CSF estimates across the early visual areas and retinal location, respectively.

**Figure 3.**
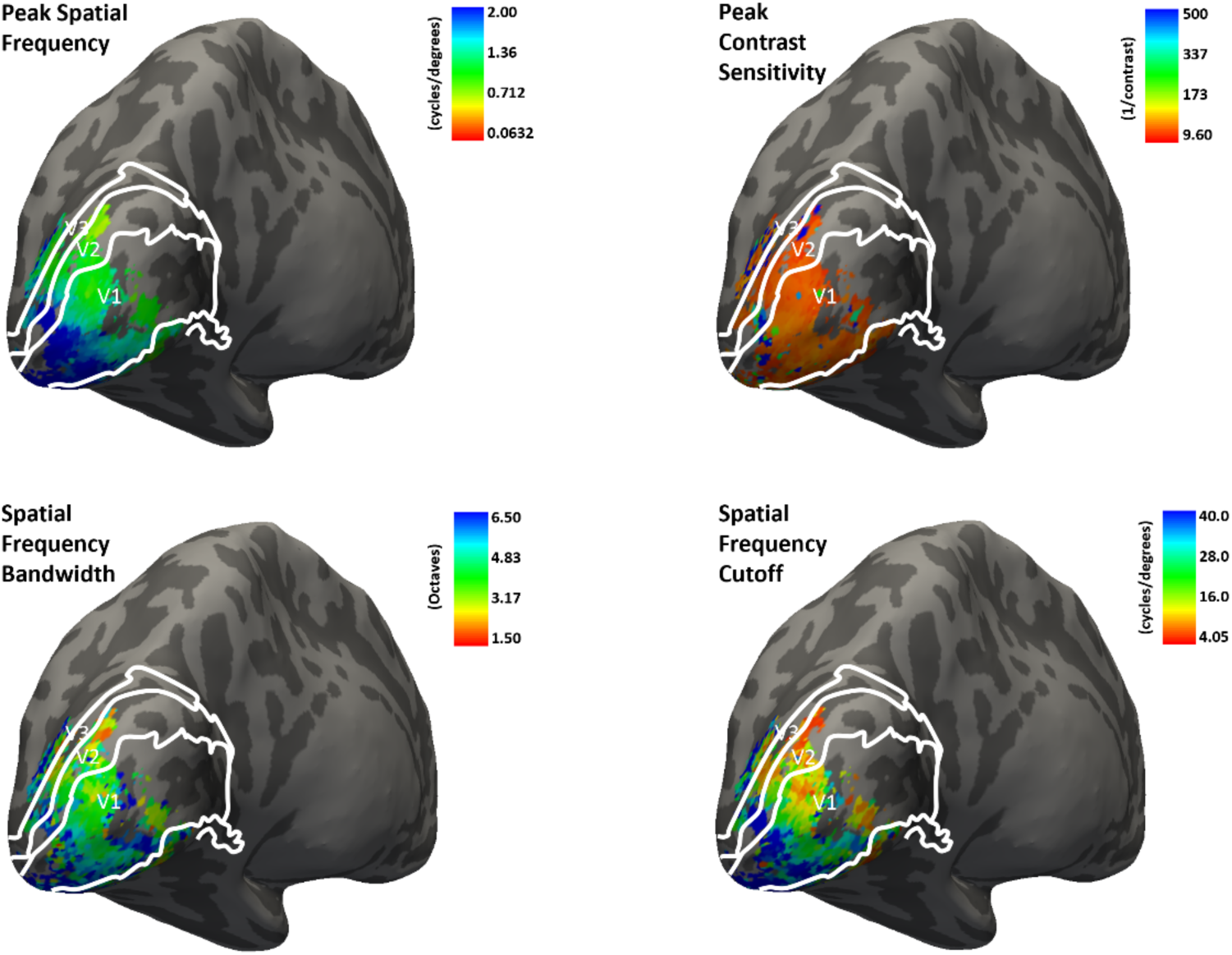
Single subject CSF parameters estimate for subject S01. Peak contrast sensitivity, peak spatial frequency, spatial frequency bandwidth, and spatial frequency cutoff estimates from 0-6° eccentricity are plotted on the inflated hemisphere of a single subject. Solid white lines indicate the borders between visual areas V1-V2-V3 obtained via N. C. Benson et al. (2012).

**Figure 4.**
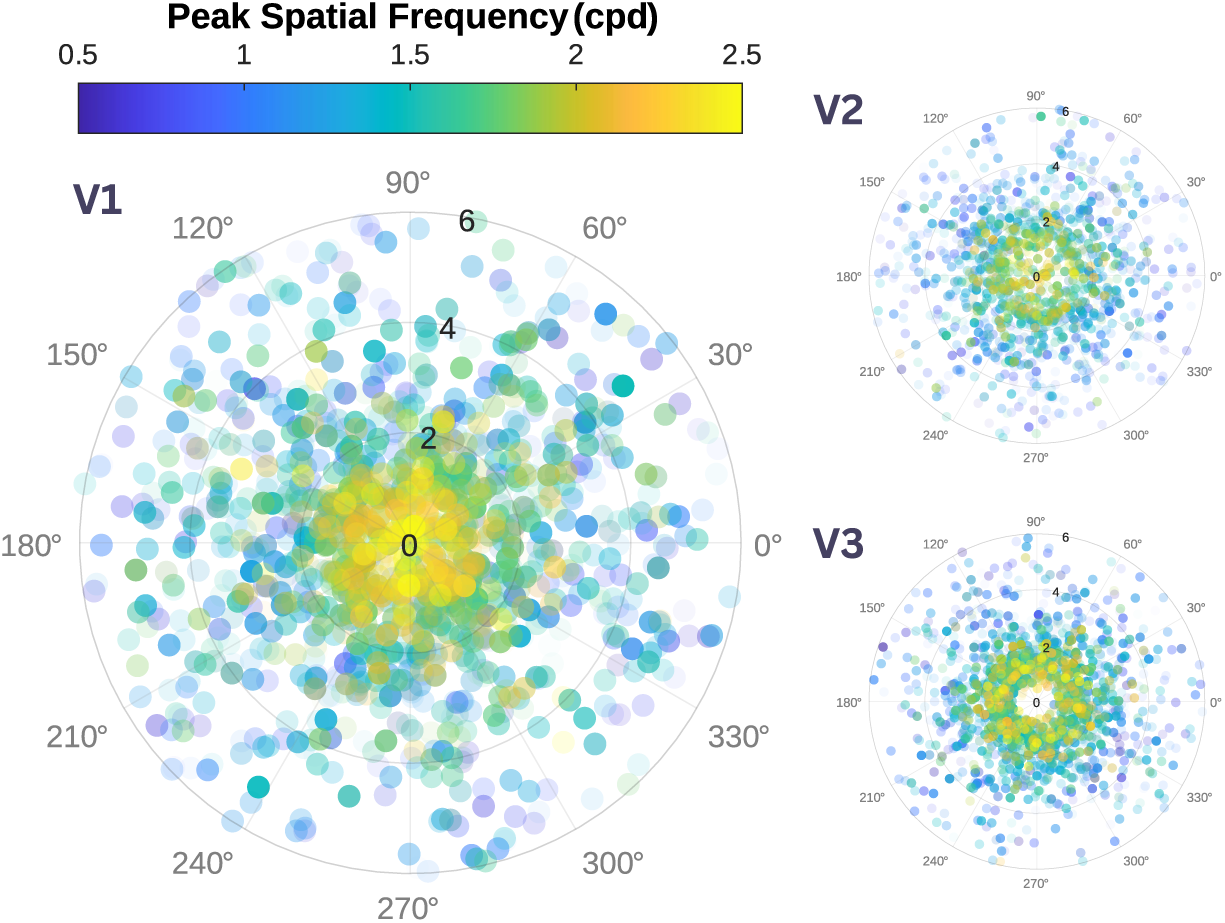
Voxel-wise peak spatial frequency map across the visual field of early visual cortex. Each circle represents a single voxel. The magnitude of the voxel’s peak spatial frequency in cycles per degree (cpd) is represented by a color code. Color Transparency of the circles are scaled with their coefficient of determination (R^2^) value for the neuroCSF fits. Their locations are determined based on their polar angle and eccentricity coordinates. In all 3 visual areas (V1, V2, V3), peak spatial frequency is higher for foveal eccentricity and decreases at parafoveal eccentricities.

For all four parameters, the trends we observed were similar across early visual areas (Fig. 3). We observed systematic changes in peak spatial frequency selectivity with eccentricity. Figure 4 represents the magnitude of the voxel’s peak spatial frequency across the visual field for areas V1, V2, V3. Consistent with previous reports (Aghajari, Vinke, & Ling, 2020; Broderick, Simoncelli, & Winawer, 2022; Henriksson, Nurminen, Hyvärinen, & Vanni, 2008; Hess, Li, Mansouri, Thompson, & Hansen, 2009), voxels with foveal retinotopic preferences were selective for higher spatial frequencies and this peak preference dropped as a function of eccentricity.

Peak sensitivity values did not show significant shifts with eccentricity across all three visual areas (Himmelberg, Winawer, & Carrasco, 2020; Rovamo, Franssila, & Näsänen, 1992; Rovamo, Virsu, & NÄSÄNen, 1978) (Fig. 5). This result aligns with previous studies, indicating that our stimuli compensated for the reduced cortical representation of peripheral retinal eccentricities (Himmelberg et al., 2020; Rovamo & Virsu, 1979; Rovamo et al., 1978; Virsu & Rovamo, 1979).

**Figure 5.**
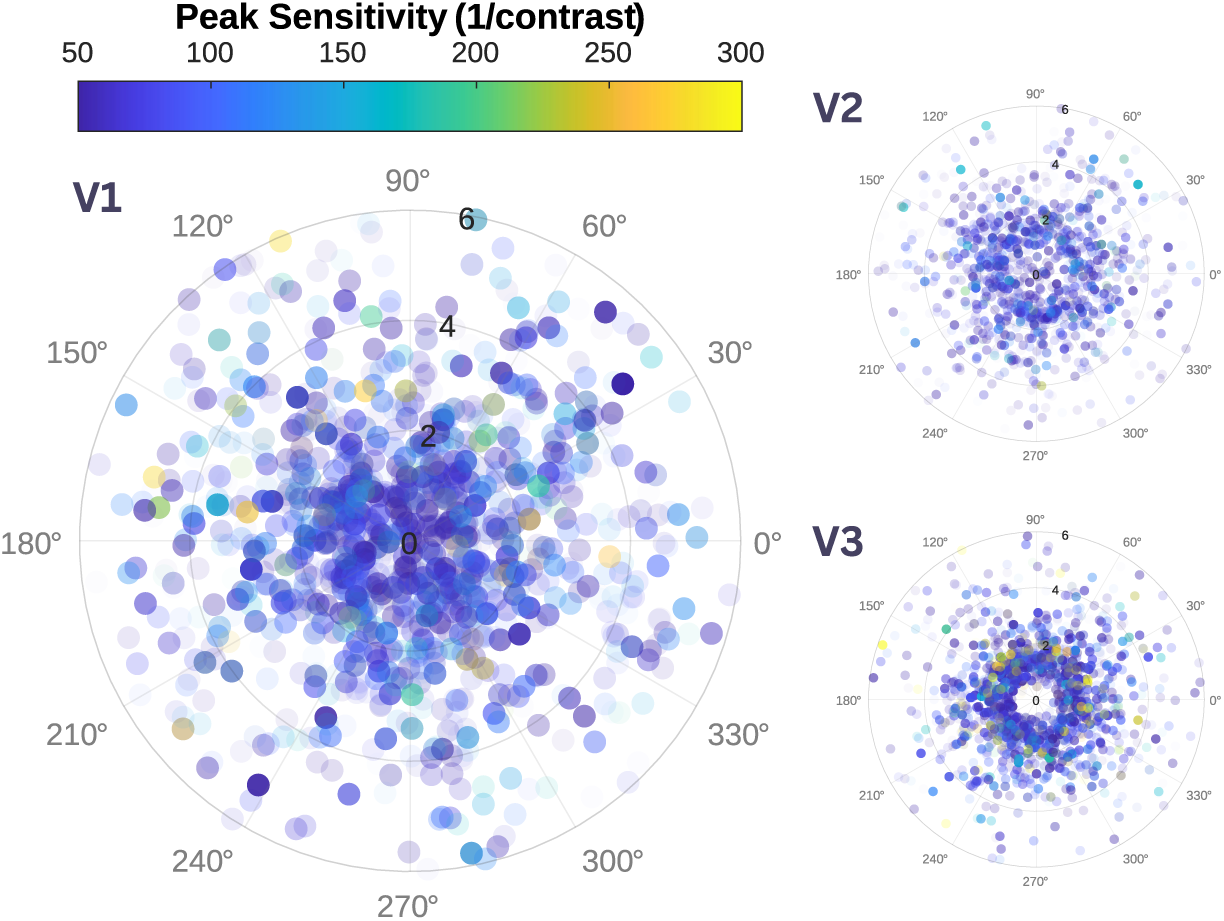
Voxel-wise peak contrast sensitivity map across the visual field of early visual cortex. Each circle represents a single voxel. The magnitude of the voxel’s peak sensitivity (1/contrast) is represented by a color code. Color Transparency of the circles are scaled with their coefficient of determination (R2) value for the neuroCSF fits. Their locations are determined based on their polar angle and eccentricity coordinates. We can see that peak sensitivity values do not shift with eccentricity across all early visual areas (V1, V2, V3).

We observed a positive trend between spatial frequency bandwidth and retinal eccentricity in V1 and V2 (Fig. 6). However, the relationship was not as clear for V3 voxels. Previous findings from human and animal studies do not predict a clear relationship between spatial frequency bandwidth and eccentricity. While Broderick et al. (2022); De Valois, Albrecht, and Thorell (1982) found that bandwidth is mostly constant across eccentricity, Aghajari et al. (2020); Foster, Gaska, Nagler, and Pollen (1985) only found a moderate increase in bandwidth with eccentricity. In both cases, that increase was more pronounced in V1 compared to higher visual areas. Our results are not inconsistent with those findings.

**Figure 6.**
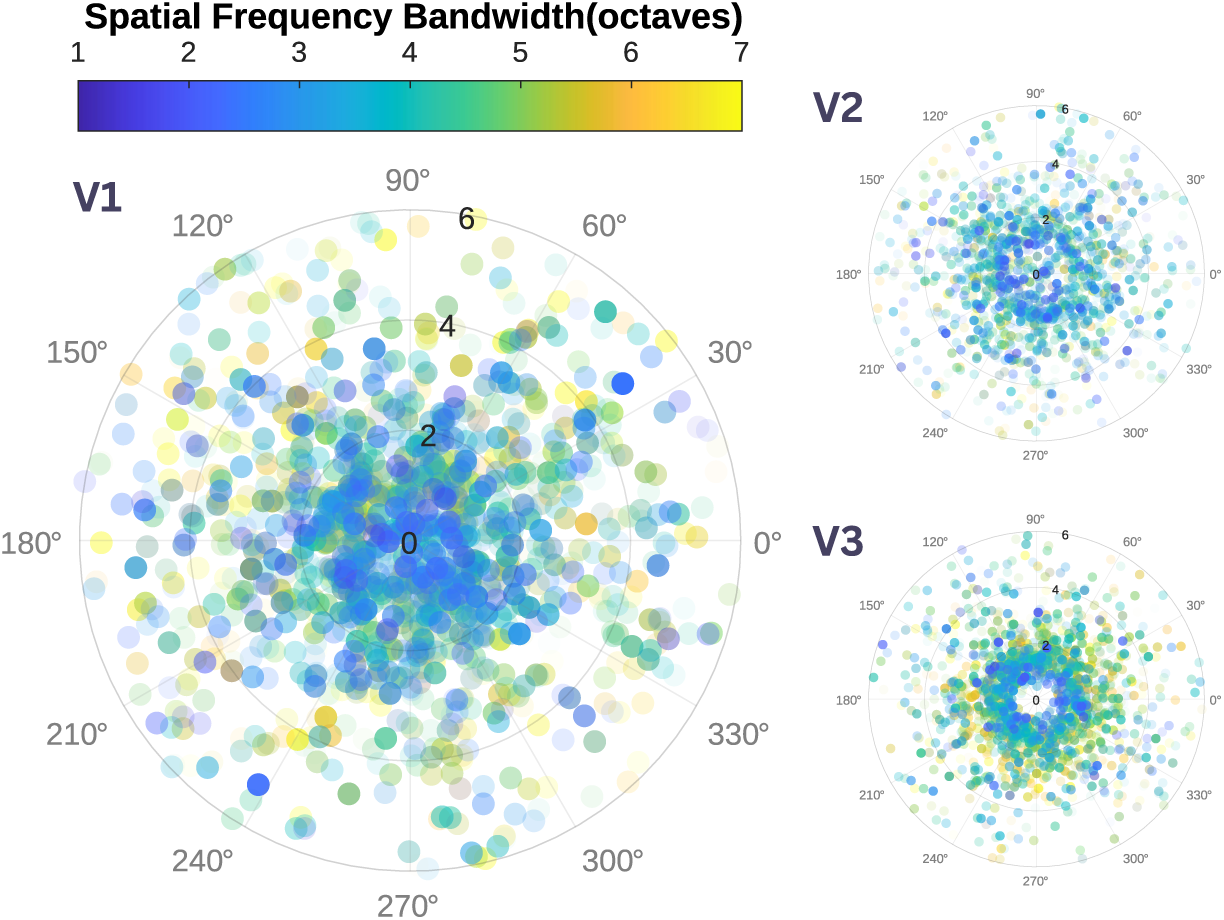
Voxel-wise spatial frequency bandwidth map across the visual field of early visual cortex. Each circle represents a single voxel. The magnitude of the voxel’s spatial frequency bandwidth (octaves) is represented by a color code. Color Transparency of the circles are scaled with their coefficient of determination (R2) value for the neuroCSF fits. Their locations are determined based on their polar angle and eccentricity coordinates. For area V1 and V2, foveal spatial frequency bandwidth is narrower compared to the peripheral region. However, there is no clear trend for V3 voxels.

Spatial frequency cutoff showed a modest drop in parafoveal regions compared to foveal regions, primarily in areas V2 and V3 (Fig. 7). High spatial frequency cutoffs remained mostly constant across eccentricities for area V1.

**Figure 7.**
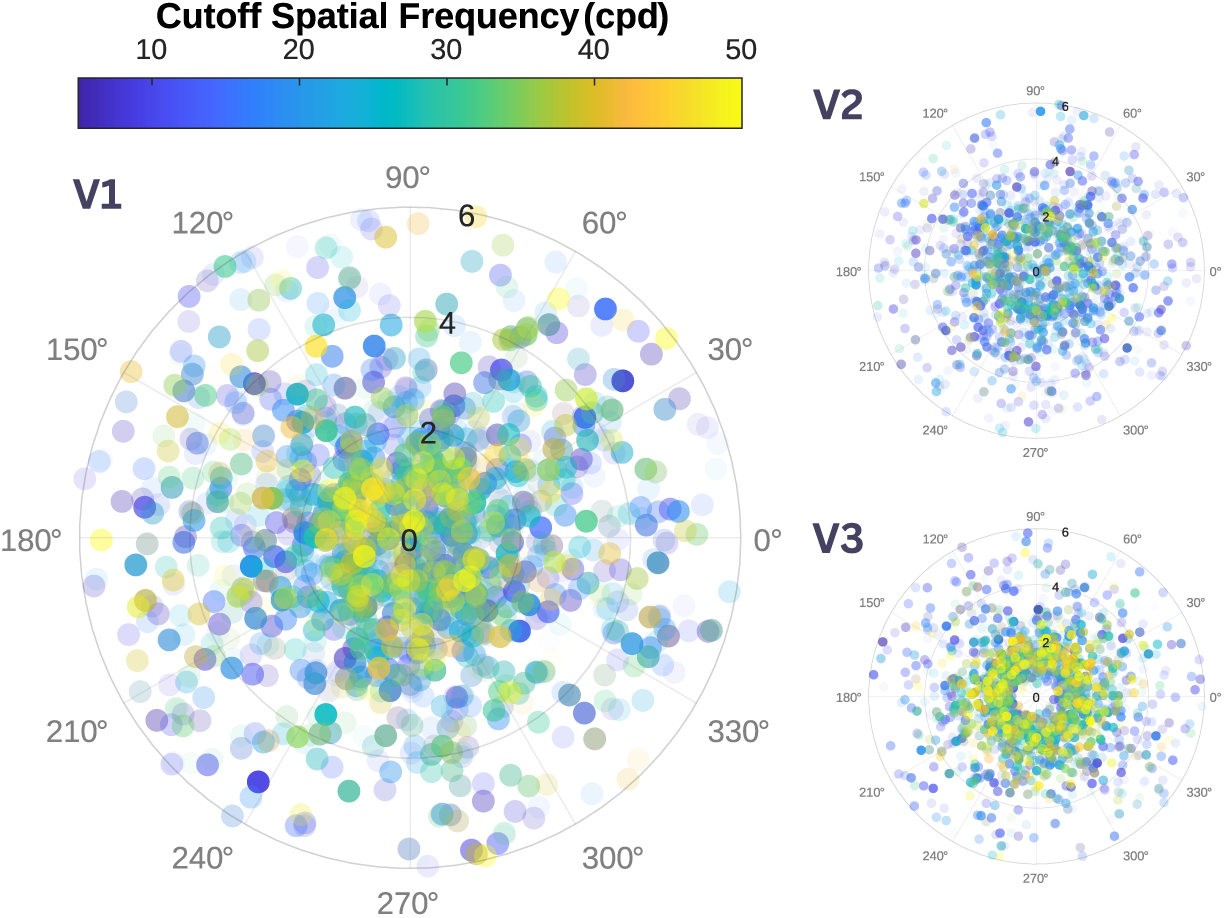
Voxel-wise spatial frequency cutoff map across the visual field of early visual cortex. Each circle represents a single voxel. The magnitude of the voxel’s spatial frequency cutoff (cpd) is represented by a color code. Color Transparency of the circles are scaled with their coefficient of determination (R2) value for the neuroCSF fits. Their locations are determined based on their polar angle and eccentricity coordinates. We cannot see a clear trend between V1 voxels spatial frequency cutoff and eccentricity, while spatial frequency cutoff values decrease with eccentricity across V2 and V3.

To focus on the eccentricity-based effects, in the next analyses we collapsed our results across polar angles, and we binned the data of our subject into 12 equal sized eccentricity bins from zero to six degrees of visual angle (dva). We looked at the average parameter response as a function of eccentricity and across early visual areas.

Previously observed trends were present across all CSF and visual areas (Fig. 8). We found a negative relationship in V1-V3 when examining the average peak spatial frequency [V1: r(12) = −0.926, P <0.001; V2: r(12) = −0.820 P <0.001; V3: r(12) = −0.792, P <0.001] and spatial frequency cutoff [V1: r(12) = −0.709, P <0.01; V2: r(12) = −0.666, P <0.01; V3: r(12) = −0.796, P <0.001] as a function of eccentricity across subjects, (Fig. 8a,d). We observed a positive trend between spatial frequency bandwidth and retinal eccentricity but only for areas V1 and V2 [V1: r(12) = 0.887, P <0.001; V2: r(12) = 0.632, P <0.05; V3: r(12) = 0.475, P = 0.086](Fig. 8c). Finally, our analysis didn’t reveal any association between peak contrast sensitivity and eccentricity [V1: r(12) =0.203, P = 0.486; V2: r(12) = 0.177, P = 0.545; V3: r(12) = 0.326, P = 0.256] (Fig. 8b).

**Figure 8.**
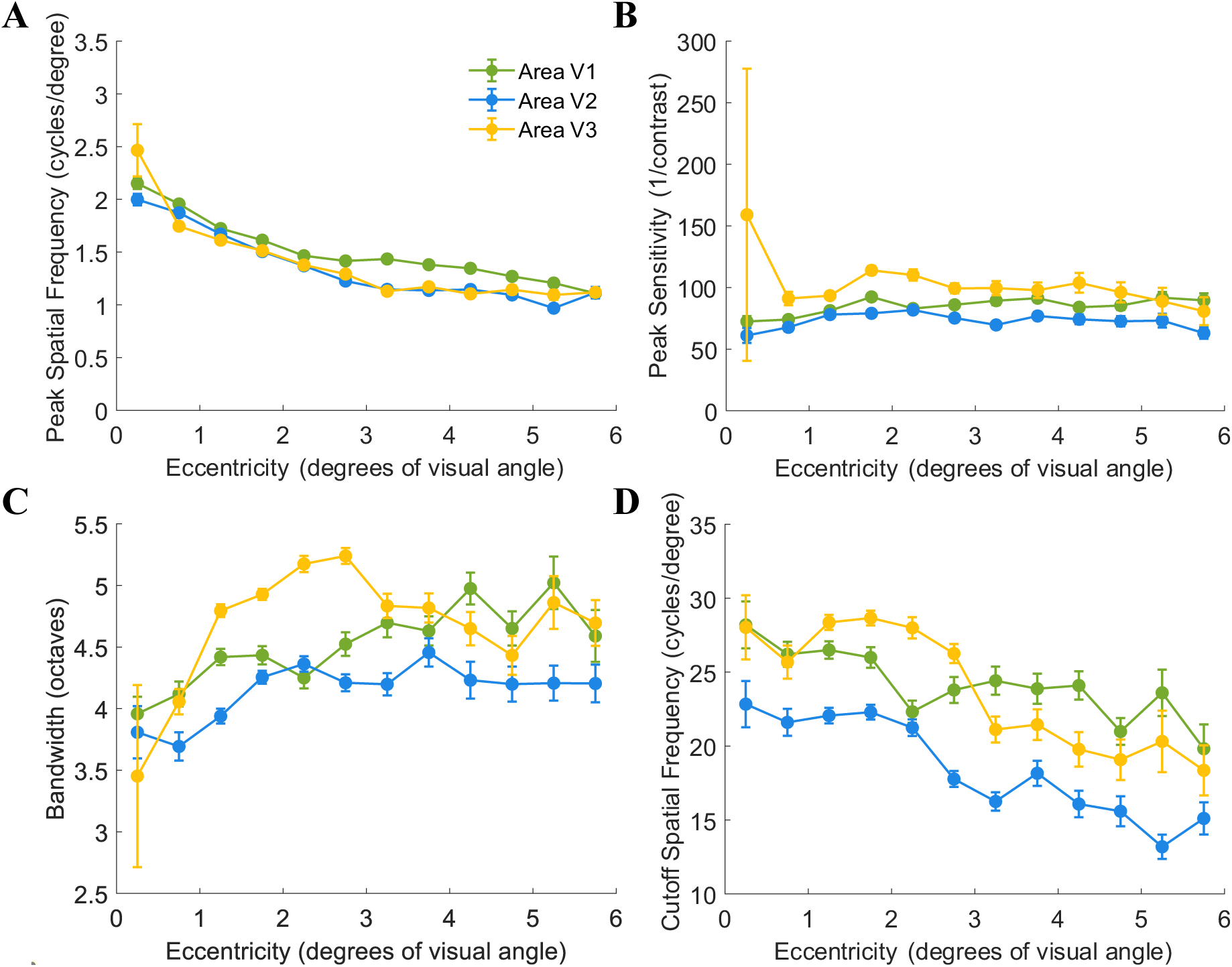
Relationship between the CSF parameters and retinotopic eccentricity across early visual areas. (A) Mean subject-wise peak spatial frequency estimates within each eccentricity bin. Peak spatial frequency is higher for foveal eccentricity and decreases at parafoveal eccentricities across V1-V2-V3. (B) Mean subject-wise peak sensitivity estimates within each eccentricity bin. Peak gain is stable across eccentricity for all three visual areas. (C) Mean subject-wise spatial frequency bandwidth estimates within each eccentricity bin. Spatial frequency bandwidth slightly increases with eccentricity in area V1 and V3. In area V2, spatial frequency bandwidth is roughly stable across eccentricity. (D) Mean subject-wise spatial frequency cutoff estimates within each eccentricity bin. In V1-V3, mean cutoff values are higher for the fovea compared to parafoveal eccentricities. 12 bins were linearly spaced between the eccentricity range [0.25°,5.75°]. Data from early visual areas (V1, V2, V3) is displayed by a different color. Error bars show the mean ± SE.

We then calculated average parameters responses for foveal (i.e., 0-2 dva) and parafoveal (i.e., 2-6 dva) eccentricities. With the help of equation 1, we defined CSF curves for foveal and parafoveal eccentricities. Each panel of figure 9a displays two mean CSF curves, one for foveal eccentricities and another for parafoveal eccentricities, for area V1-V2-V3 for all our subjects. We then averaged these parameters to create averaged CSF curves across all our subjects and performed a Wilcoxon Signed-Rank test (Fig. 9b). The prototypical behavioral CSF curve from Lesmes, Lu, Baek, and Albright (2010) is also included as a reference.

**Figure 9.**
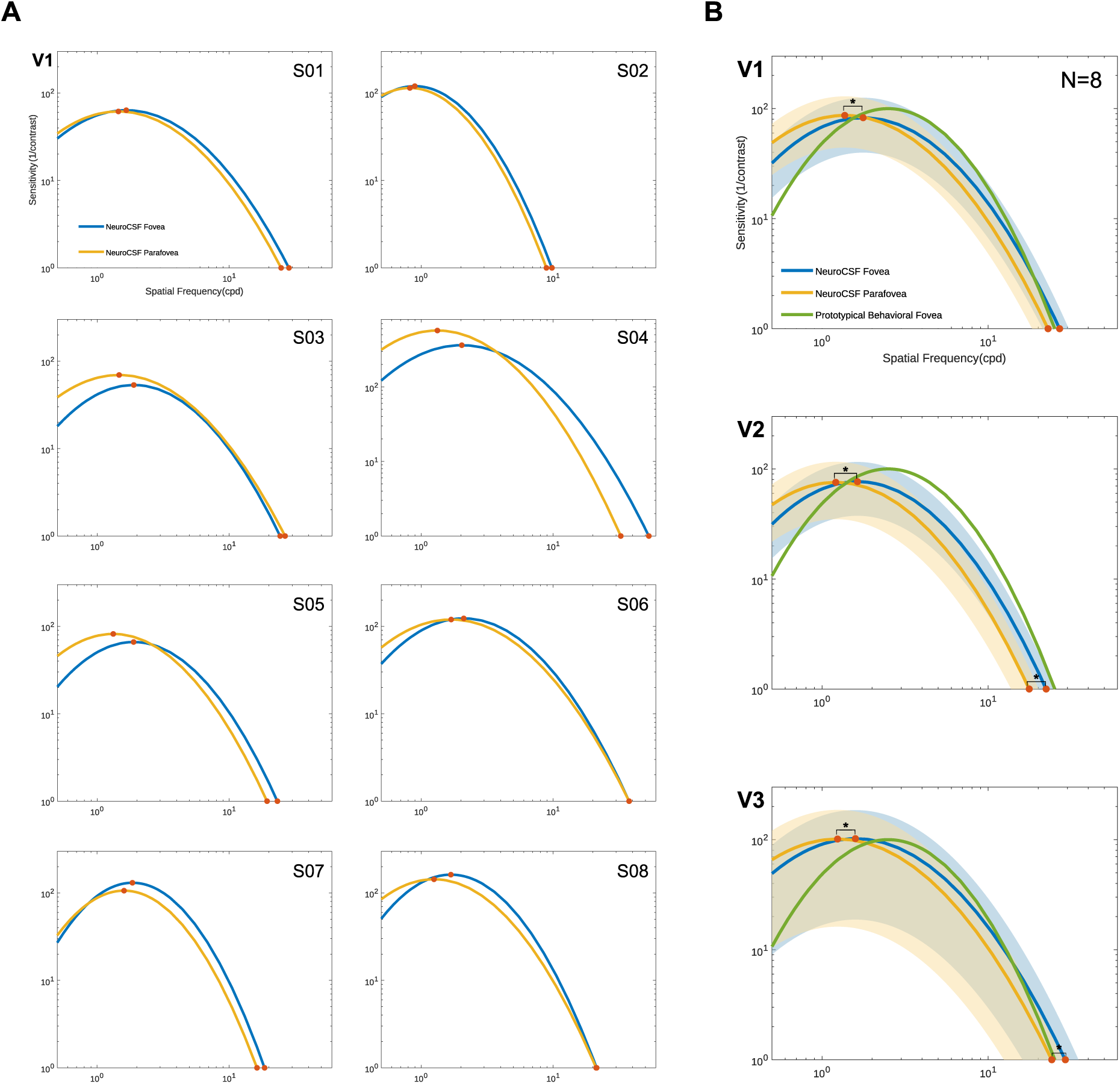
Estimated CSFs across early visual areas. (A) Estimated individualized CSFs across V1. Each panel displays one set of estimated CSFs for area V1 of one of our 8 subjects. (B) Average ± standard deviation of the CSF across areas V1, V2, and V3. Data from foveal (0-2 dva) and parafoveal (2-6 dva) eccentricities is displayed by a different color, blue and yellow respectively. The prototypical behavioral CSF curve from Lesmes et al. (2010) is also included in green as a reference. The shading represents one standard deviation above and below the mean. Significant differences (P < 0.05) between neuroCSF fovea and parafovea are indicated by a (*).

The results show that there is a statistically significant difference in the mean response of peak spatial frequency between foveal and parafoveal eccentricities in V1-V3 [V1: *Z = −2.660*, *P < 0.01*; V2: *Z = −2.660*, *P < 0.01*; V3: *Z = −2.660*, *P < 0.01*], as well as spatial frequency cutoff in V2-V3 [V1: *Z = −1.295*, *P = 0.195*; V2: *Z = −2.266*, *P < 0.05*; V3: *Z = −2.266*, *P < 0.05*]. However, there were no statistically significant differences in peak contrast sensitivity [V1: *Z = - 0.069*, *P = 0.945*; V2: *Z = −0.197*, *P = 0.844*; V3: *Z = −0.329*, *P = 0.742*] and spatial frequency bandwidth [V1: *Z = 1.762*, *P = 0.078*; V2: *Z = 0.873*, *P = 0.383*; V3: *Z = 1.762*, *P = 0.078*] between fovea and parafovea.

### 3.2. Intraclass Correlation Coefficient (ICC) Analysis for Reliability Assessment

In many settings, including clinical research, reducing the number of acquisitions can significantly decrease participant burden, scanner time, and associated costs. We performed an analysis to optimize future data collection, balancing reliability with time and cost efficiency. The goal was to identify the minimal number of runs needed to maintain data quality equivalent to that obtained with the maximal number of runs in this experiment.

To evaluate the reliability of our measurements across different run combinations, we performed an Intraclass Correlation Coefficient (ICC) analysis. The ICC is a widely used reliability index that quantifies both the degree of correlation and agreement among repeated measurements. We aimed to determine the minimal number of runs necessary to achieve reliability comparable to our gold-standard while minimizing time and financial costs associated with data collection. Our gold-standard for reliability was defined as the ICC value obtained by averaging across three identical pairs of runs, which represents the highest achievable level of consistency we could obtain from the data collected and serves as the benchmark against which all other run combinations were compared. To this end, we calculated ICC values for a range of different run combinations, including: one individual run, two individual runs, three individual runs, four individual runs, five individual runs, six individual runs, one identical pair of runs and two identical pairs of runs (Table 1). Individual runs refer to runs processed independently to generate unique sets of CSF parameters, which are then averaged to assess reliability. In contrast, pairs of identical runs involve averaging the timeseries of sets of runs with the same stimuli before processing to yield a single set of CSF parameters for each pair.

**Table 1.**
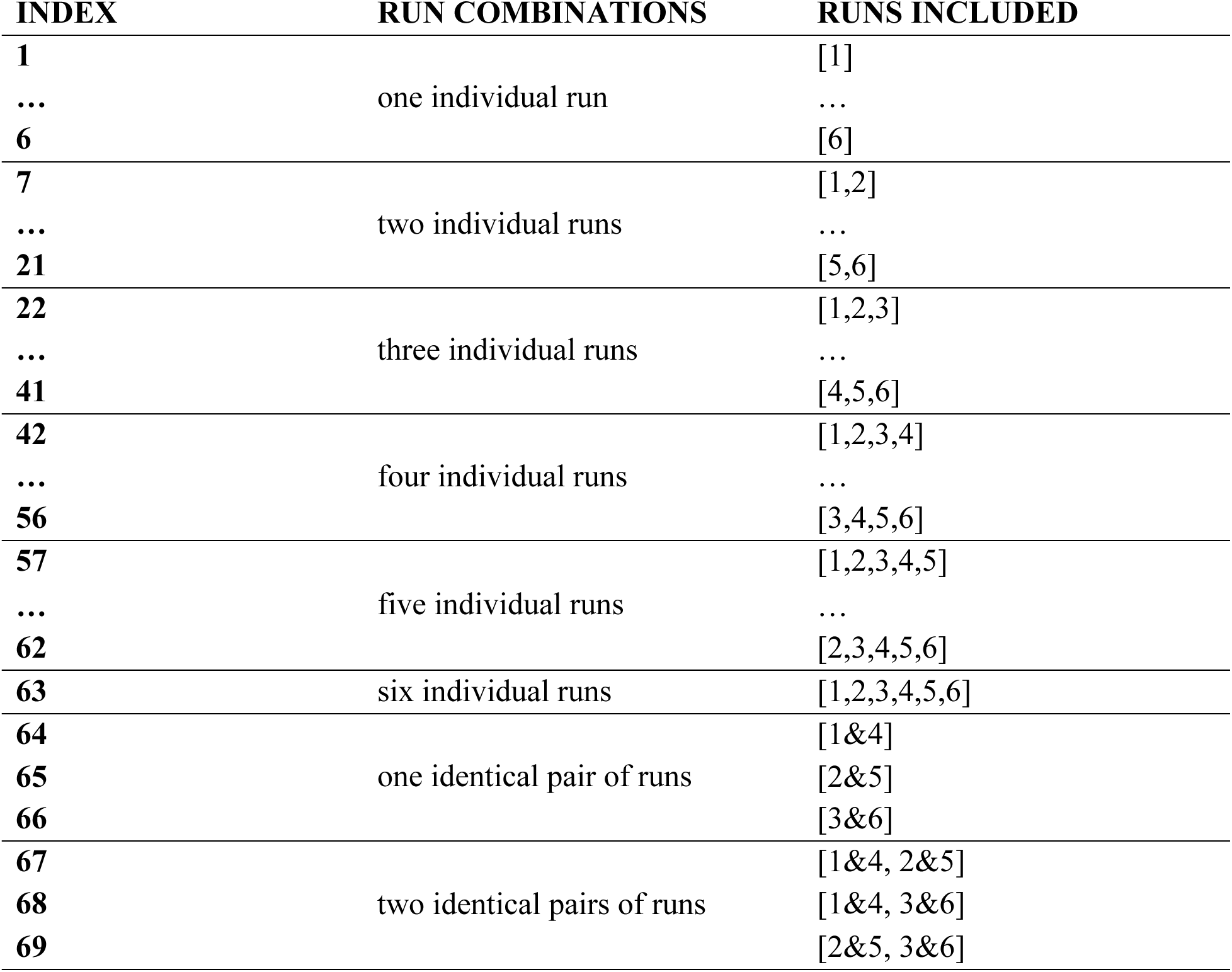
Short List of the 69 Run Combinations Included in the Intraclass Correlation Coefficient (ICC) Analysis. The table details a selection of the combinations of runs used in the ICC analysis, organized by the number of runs included: from single individual runs to multiple individual runs, as well as pairs of identical runs that were averaged together. The symbol "," denotes combinations of runs whose time series were individually processed to generate separate sets of contrast sensitivity function (CSF) parameters, which were then averaged. The symbol "&" represents combinations of runs with identical sets of stimuli, where the time series were averaged prior to processing to yield a single set of CSF parameters.

| INDEX | RUN COMBINATIONS | RUNS INCLUDED |
| --- | --- | --- |
| 1 |  | [1] |
| ... | one individual run | ... |
| 6 |  | [6] |
| 7 |  | [1,2] |
| ... | two individual runs | ... |
| 21 |  | [5,6] |
| 22 |  | [1,2,3] |
| ... | three individual runs | ... |
| 41 |  | [4,5,6] |
| 42 |  | [1,2,3,4] |
| ... | four individual runs | ... |
| 56 |  | [3,4,5,6] |
| 57 |  | [1,2,3,4,5] |
| ... | five individual runs | ... |
| 62 |  | [2,3,4,5,6] |
| 63 | six individual runs | [1,2,3,4,5,6] |
| 64 |  | [1&4] |
| 65 | one identical pair of runs | [2&5] |
| 66 |  | [3&6] |
| 67 |  | [1&4, 2&5] |
| 68 | two identical pairs of runs | [1&4, 3&6] |
| 69 |  | [2&5, 3&6] |

To determine acceptable reliability, we selected an ICC threshold of 0.5 (Koo & Li, 2016; Liljequist, Elfving, & Skavberg Roaldsen, 2019). An ICC value below 0.5 was considered to indicate poor reliability. The aim was to determine which of these combinations could achieve reliability equivalent to or greater than our gold-standard, thereby providing a reliable measure while using the fewest possible runs.

A one-sample t-test was conducted to compare the ICC values of the eight different run combinations against the threshold value of 0.5 across five parameters: peak contrast sensitivity, peak spatial frequency, spatial frequency bandwidth, spatial frequency cutoff, and R² value. The statistical significance of the results was evaluated using p-values, with Bonferroni correction applied for multiple comparisons to control for Type I error. Run combinations with p-values below 0.05 after correction were considered statistically significant.

Out of the eight combinations tested, only "one identical pair of runs” and "two identical pairs of runs" achieved reliability greater than our threshold value (Fig. 10). The results were similar across all parameters. For "one identical pair of runs," significant results were found for peak contrast sensitivity (t(7) = 5.870, p <0.01), peak spatial frequency (t(7) = 5.513, p < 0.01), spatial frequency bandwidth (t(7) = 6.075, p < 0.01), spatial frequency cutoff (t(7) = 12.703, p < 0.001), and R² value (t(7) = 12.970, p < 0.001). The "two identical pairs of runs" also showed significant results for peak contrast sensitivity (t(7) = 15.581, p < 0.001), peak spatial frequency (t(7) = 10.051, p < 0.001), spatial frequency bandwidth (t(7) = 16.371, p < 0.001), spatial frequency cutoff (t(7) = 32.722, p < 0.001), and R² value (t(7) = 27.398, p < 0.001). None of the other run combinations yielded results that were above the threshold across any of the parameters tested (all p > 0.05). These findings suggest that it is possible to reduce the number of runs and still maintain robust data quality, offering a more efficient and economical approach to data collection.

**Figure 10.**
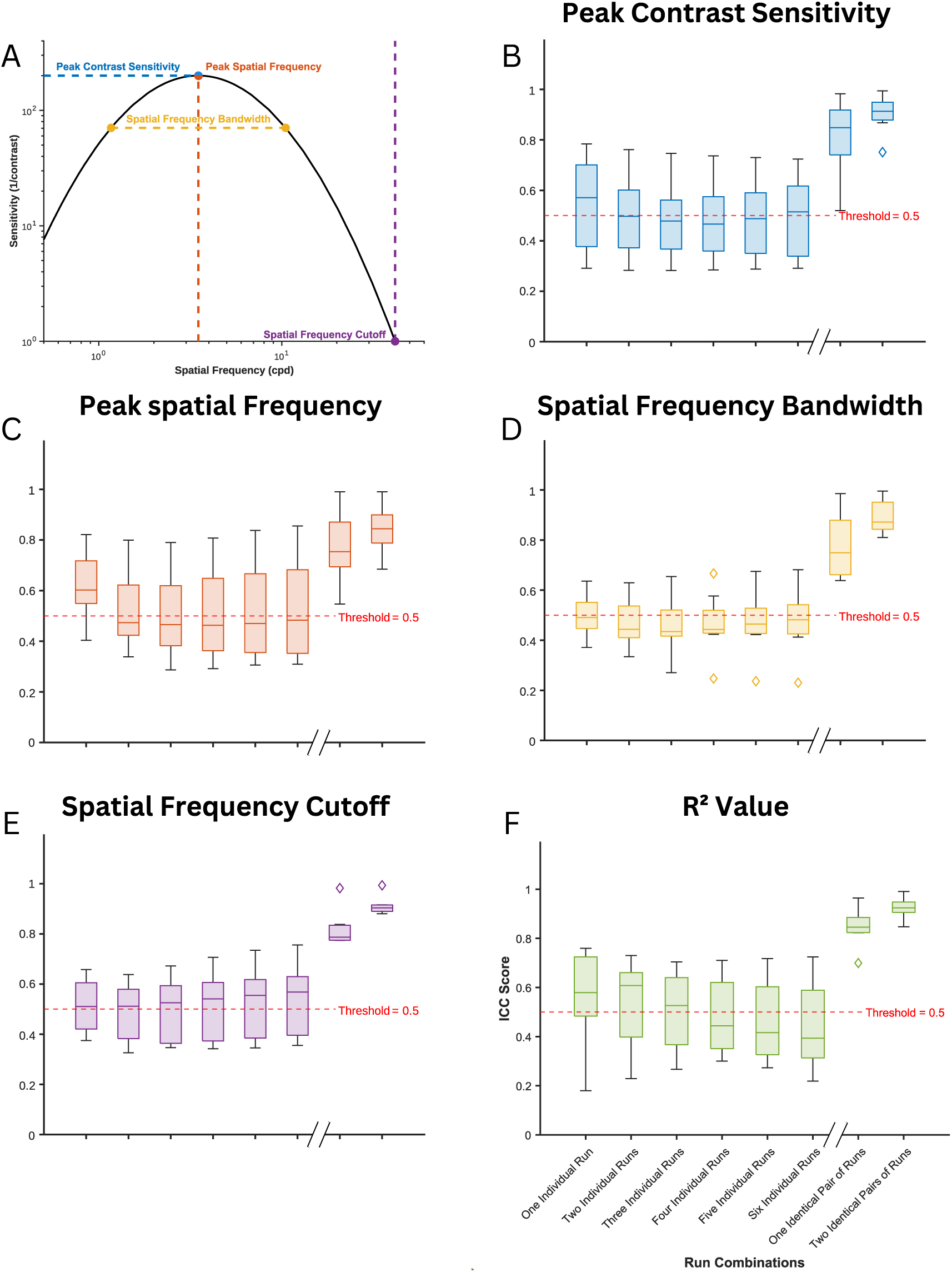
Distribution of Intraclass Correlation Coefficient (ICC) values for all combinations of runs compared to the gold standard across five CSF parameters. (A) The Contrast Sensitivity Function (CSF) curve with markers indicating the four parameters: peak contrast sensitivity (blue), peak spatial frequency (red), spatial frequency bandwidth (orange), and spatial frequency cutoff (purple). (B-F) ICC values across different run combinations for each parameter: (B) Peak Contrast Sensitivity, (C) Peak Spatial Frequency, (D) Spatial Frequency Bandwidth, (E) Spatial Frequency Cutoff, and (F) R² Value. The horizontal red dashed line at ICC = 0.5 indicates the threshold for acceptable reliability. Asterisks (*) denote run combinations that are statistically significantly different from the threshold (ICC score=0.5).

## 4. Discussion

The neuroCSF is a novel fMRI model-driven approach that combines dynamic stimulation, with a canonical model of the CSF and parameter optimization to provide voxel-wise CSF estimations across early visual areas. With this new method, we were able to derive robust and interpretable estimates of cortical CSF parameters to provide the first complete characterization of the CSF using neuroimaging data. Our approach provides estimates of four parameters per visual field location: peak gain, peak spatial frequency, spatial frequency bandwidth, and the cut-off frequency of the CSF.

One of the major concerns of vision neuroscience is to relate the organization of cortical processing to perception. Psychophysical studies have consistently shown that behavioral measures of contrast and spatial frequency sensitivities decrease from foveal to peripheral eccentricities. Previous neuroimaging and neurophysiological studies independently estimated human spatial frequency tuning (Aghajari et al., 2020; Broderick et al., 2022; Henriksson et al., 2008; Singh, Smith, & Greenlee, 2000; Sirovich & Uglesich, 2004) and contrast sensitivity (Himmelberg & Wade, 2019; Marquardt, Schneider, Gulban, Ivanov, & Uludağ, 2018) across the visual cortex. To our knowledge, the neuroCSF is the first human neuroimaging approach that unifies these measurements. This integration allowed us to explore the spatial organization of the CSF across early visual areas to provide valuable insights into the relationship between behavioral CSF measurements and their cortical counterparts.

The dependency of CSF parameters on retinotopy and eccentricity that we found in all three visual areas was consistent with prior studies (Aghajari et al., 2020; Broderick et al., 2022; Campbell, Cooper, & Enroth-Cugell, 1969; De Valois et al., 1982; Foster et al., 1985; Movshon, Thompson, & Tolhurst, 1978). Our results support Aghajari et al. (2020); Campbell et al. (1969); Movshon et al. (1978) findings, suggesting a fast rate of peak spatial frequency decline with eccentricity across V1-V3. Similarly to Henriksson et al. (2008), V2 voxels optimal spatial frequencies were on average 2/3 of the peak spatial frequency in V1. We also observed a similar shift from V2 to V3. Spatial frequency bandwidth changes with eccentricity were similar across V1&V3, with larger values in peripheral eccentricity. Spatial frequency bandwidth remained relatively consistent across eccentricity for V2. These results match previous conflicting reports of the relationship between spatial frequency selectivity and eccentricity (Aghajari et al., 2020; Broderick et al., 2022; De Valois et al., 1982; Foster et al., 1985). Aghajari et al. (2020) noted a slight increase in bandwidth with eccentricity but only for the perifovea. Since we only sampled the fovea and parafovea this could explain why we didn’t detect a similar trend between spatial frequency selectivity and eccentricity. These results can help resolve some of the discrepancies in the literature regarding spatial frequency selectivity and eccentricity.

Peak gain was found to be independent of eccentricity across V1-V3. By using full field stimuli, we ensured the cortical representation of the gratings at various eccentricities became equal in size, compensating for the reduced cortical representation that usually limits contrast sensitivity in the periphery (Himmelberg et al., 2020; Horton & Hoyt, 1991; Rovamo & Virsu, 1979; Rovamo et al., 1978; Virsu & Rovamo, 1979). Similarly to Broderick et al. (2022), our results did not reveal a significant trend between spatial frequency cutoff and eccentricity for area V1. When looking at the individual CSFs (Fig. 9a), we can also see that for some subjects (S03, S06, S08), the parafoveal spatial frequency cutoff was higher than the foveal one. These results could be explained by the variability in foveal size that exists across the population (Noah C. Benson et al., 2022; Henriksson, Karvonen, Salminen-Vaparanta, Railo, & Vanni, 2012; Schira, Tyler, Breakspear, & Spehar, 2009). Our estimates of retinal location were obtained from a probabilistic atlas (N. C. Benson et al., 2012) based on cortical-anatomical landmarks averaged across 25 individuals. Noah C. Benson et al. (2022) showed that the use of automated anatomical templates was less effective at estimating spatial layout of the visual field than mapping the retinotopic organization. They showed a two-fold variation in amount of cortex devoted to the fovea across individuals. This suggests that standardized atlases may not capture the true extent of variability found in individuals retinotopic maps, and hence mistakenly label parafoveal voxels as foveal ones and vice-versa. These results could also explain why we did not find a statistically significant difference between fovea and parafovea spatial frequency cutoffs in V1. However, we did find a significant effect of eccentricity on the cutoff for V2-V3 voxels.

In comparison with behavioral measures, the neuroCSF CSF curves were of similar shapes, but they were shifted along the x-axis toward lower spatial frequencies. When measured psychophysically, both human and macaque contrast sensitivity tends to peak between 3-5 (cpd) (De Valois, Morgan, & Snodderly, 1974; Watson & Ahumada, 2005), and these values shift toward lower frequencies with increasing eccentricity. Our results indicated an overall lower resolution of early visual areas voxels with mean foveal spatial frequency optima of 1.9 cpd, 1.7 cpd and 1.6 cpd for V1, V2 and V3 respectively. Comparison between cortical and behavioral results is not straightforward and several factors inherent to the differences between fMRI and psychophysical measurements of visual function could explain these discrepancies.

NeuroCSF directly measures cortical sensitivity in relation to visual stimuli, capturing neural processing in early visual cortex. This cortical activity might not have a one-to-one correspondence to perceived sensitivity. Behavioral CSF involves higher-level visual processing and cognitive compensations such as attention (Gandhi, Heeger, & Boynton, 1999; Ress, Backus, & Heeger, 2000; Ress & Heeger, 2003) and decision-making processes (e.g., learning, experience, and cognitive strategies), potentially shifting sensitivity. These cognitive processes can integrate information over time and space, which may lead to increased sensitivity, especially at higher spatial frequencies. Uncertainty related to the precise timing and perception of our stimulus might have also influenced cortical activity level. Subjects were presented with a low-contrast target that was sometimes reported as “hard to perceive” and they were allowed to completely disengage their attention during the interstimulus intervals. Ress et al. (2000) showed that fMRI measurements of cortical activity in early visual areas during a contrast detection task highly depend on attention-related signals. Our low sensitivity results might therefore be a consequence of low visual attention.

fMRI inherently has limitations in spatial resolution, and the neuroCSF results are contingent on the spatial precision of the imaging. Fine-grained CSF changes may be challenging to detect due to the limited spatial resolution of fMRI. This limitation can potentially lead to underestimations or misinterpretations of CSF parameters, especially when investigating subtle changes in visual perception. A large voxel size may result in spatial averaging of neural responses, potentially leading to a shift in the peak spatial frequency toward lower values. Therefore, careful consideration of spatial resolution constraints is crucial.

Our approach differs from existing methods to measure human CSF in that it overcomes measurement trade-offs and patient limitations, while also presenting pathophysiological relevance. Traditional approaches to measuring the human CSF face trade-offs between precision and testing time, limiting their utility. When sampling the two-dimensional space of possible stimuli-defined by contrast and spatial frequency- the range of stimuli needs to be wide enough to capture the global shape of the CSF, but it also needs to be precise enough to capture the highly dynamic regions of the curve such as the high-frequency cutoff. To improve the flexibility and precision of traditional CSF tests, sampling range needs to be increased, but this would introduce the cost of extra testing time. In contrast, with our model-driven approach that is combined with dynamic stimuli, we can sweep through a wide range of spatial frequency and contrast across the visual field over a short period of time. This strategy allows for a wide sampling range and resolution without increased testing time.

Current methods to measure the CSF can be challenging for healthy participants, and even more so for patients who suffer from attention or fixation problems and visual discomfort (Kalia et al., 2014; Lesmes et al., 2010). These factors can also influence the reliability of current behavioral CSF measures (Abrahamyan et al., 2016; Abramov et al., 1984; Bradley & Freeman, 1982; Lee et al., 2014). With its use of a full field stimuli, neuroCSF eliminates the need for participant fixation. Our novel technique also estimates the CSF without the need for participant input, making it suitable for nonverbal individuals like infants or those with intellectual disabilities. While neuroCSF offers advantages in stimulus flexibility, optimizing stimulus designs to accurately capture the complete CSF profile is not without challenges. Balancing the range and resolution of stimuli while minimizing testing time remains a methodological concern that needs careful consideration. Next iterations of the stimulus could be more targeted to reduce the scanning time. We are currently working on an optimization approach to define a set of stimuli that yields the most information about the CSF’s overall shape.

The use of ICC analysis helped us identify the optimal number and types of runs required to achieve high reliability with neuroCSF measurements. The results suggest that while the use of a single or multiple individual runs often yields moderate ICC values (i.e., below the threshold of 0.5), combining identical runs before data processing significantly enhances reliability, exceeding the 0.5 threshold for ICC scores in all cases. By identifying combinations of runs that can achieve reliable measurements with fewer repetitions, our analysis offers a practical framework for optimizing data collection. This has important implications for future studies, where minimizing participant burden and resource usage is crucial.

Although the neuroCSF technique provides valuable insights into cortical CSF parameters, its relationship to traditional behavioral CSF measures should be further validated. Understanding the degree of correspondence between neuroimaging-derived CSF estimates and psychophysical measurements is crucial for establishing its reliability and validity.

The neuroCSF technique relies on a log-parabolic model to describe CSF parameters. This model of the CSF is a simplified version of the truncated log-parabola that has previously been used in psychophysical studies to estimate CSF parameters (Lesmes et al., 2010; Rosenkranz et al., 2021; Tardif, Watson, Giaschi, & Gosselin, 2021; Watson & Ahumada, 2005), and has been shown to provide, compared to other models, excellent fits with very few assumptions, parameters and calculations (Gao et al., 2015; Reynaud, Tang, Zhou, & Hess, 2014; Spiegel et al., 2016; Watson & Ahumada, 2005). While this model simplifies the estimation process, it may not fully capture all the nuances of individual differences or variations in CSF profiles. The symmetric log-parabola typically misfits the plateau observed on the peak’s low-frequency side (Rohaly & Owsley, 1993). This could constrain the model’s accuracy in specific cases. With an additional parameter to describe the low-frequency plateau, the truncated log-parabola can deal with the issues of the CSF’s asymmetry.

Addressing these limitations and conducting further research to mitigate their impact will be essential for harnessing the full potential of the neuroCSF technique.

## 5. Data and Code Availability

The data that support the findings of this study are available on request from the corresponding author.

## 6. Authors Contributions

**Laurie Goulet:** Conceptualization, Methodology, Software, Formal analysis, Investigation, Writing – Original Draft, Visualization. **Reza Farivar:** Conceptualization, Methodology, Resources, Writing – Review & Editing, Project administration, Funding acquisition.

## 7. Acknowledgements

This research was partly supported by CIHR under Grant Number 899956. The authors declare no competing interests.

